# Reconstructing the human first trimester fetal–maternal interface using single cell transcriptomics

**DOI:** 10.1101/429589

**Authors:** Roser Vento-Tormo, Mirjana Efremova, Rachel A. Botting, Margherita Y. Turco, Miquel Vento-Tormo, Kerstin B. Meyer, Jongeun Park, Emily Stephenson, Krzysztof Polański, Rebecca P. Payne, Angela Goncalves, Angela Zou, Johan Henriksson, Laura Wood, Steve Lisgo, Andrew Filby, Gavin J. Wright, Michael J. T. Stubbington, Muzlifah Haniffa, Ashley Moffett, Sarah A. Teichmann

**Affiliations:** Wellcome Trust Sanger Institute, Wellcome Genome Campus, Hinxton, Cambridge CB10 1SA, United Kingdom; Institute of Cellular Medicine, Newcastle University, Newcastle upon Tyne, NE2 4HH, UK; Department of Pathology, University of Cambridge, Cambridge, CB2 1QP, UK; Centre for Trophoblast Research, University of Cambridge, UK; Department of Physiology, Development and Neuroscience, University of Cambridge, Cambridge, CB2 1QP, UK; YDEVS software development, Valencia 46009, Spain; German Cancer Research Center (DKFZ), Heidelberg, Germany; Human Developmental Biology Resource, Institute of Genetic Medicine, Newcastle University, Newcastle upon Tyne, NE1 3BZ; Department of Dermatology and NIHR Newcastle Biomedical Research Centre, Newcastle Hospitals NHS Foundation Trust, Newcastle upon Tyne NE2 4LP, UK; Theory of Condensed Matter Group, The Cavendish Laboratory, University of Cambridge, Cambridge, UK; European Molecular Biology Laboratory, European Bioinformatics Institute (EMBLEBI), Hinxton, Cambridge, UK

## Abstract

During the early weeks of human pregnancy, the fetal placenta implants into the uterine mucosa (decidua) where placental trophoblast cells intermingle and communicate with maternal cells. Here, we profile transcriptomes of ∼50,000 single cells from this unique microenvironment, sampling matched first trimester maternal blood and decidua, and fetal cells from the placenta itself. We define the cellular composition of human decidua, revealing five distinct subsets of decidual fibroblasts with differing growth factors and hormone production profiles, and show that fibroblast states define two distinct decidual layers. Among decidual NK cells, we resolve three subsets, each with a different immunomodulatory and chemokine profile. We develop a repository of ligand-receptor pairs (www.CellPhoneDB.org) and a statistical tool to predict the probability of cell-cell interactions *via* these pairs, highlighting specific interactions between decidual NK cells and invading fetal extravillous trophoblast cells, maternal immune and stromal cells. Our single cell atlas of the maternal-fetal interface reveals the cellular organization and interactions critical for placentation and reproductive success.

---

During early pregnancy, the placenta implants into the uterine mucosal lining, the decidua. Normal formation of the placenta and access to the maternal arterial blood supply determine reproductive success. Indeed, major complications of pregnancy such as pre-eclampsia and fetal growth restriction have their origins in very early pregnancy^1^. During pregnancy, the endometrium is transformed to decidua under the influence of progesterone secreted by the *corpus luteum*. Decidualisation results from a complex but well-orchestrated cellular differentiation program including enlargement of the stromal cells, increase in secretory activity of the glandular cells, and the expansion of the dominant immune population in the decidua, the distinctive decidual NK cells (dNK)^2^.

Following implantation of the blastocyst through the surface epithelium, placental trophoblast cells invade through the decidua and move towards the spiral arteries, where they destroy the smooth muscle media to transform the arteries into high conductance vessels^3^. Extravillous trophoblast (EVT) migrates right through the decidua and reaches as far as the inner myometrium. Balanced regulation of EVT invasion is critical to pregnancy success: arteries must be sufficiently transformed, but excessive invasion prevented, to ensure correct allocation of resources to both mother and baby. The decidua plays a key role in regulating invasion and when it is absent, for example when the placenta implants on a previous cesarean section scar, trophoblast invades uncontrollably, threatening the viability of the pregnancy and maternal wellbeing.^4,5^.

The different cellular components of decidua all support the establishment of early pregnancy in human. Decidual glands provide histiotrophic nutrition to the placenta before the arterial connections are established^6,7^. Stromal cells generate a rich extracellular matrix (ECM), that facilitates EVT invasion, and secrete products such as prolactin and insulin-like growth factor-binding protein 1 (IGFBP-1)^2^. Damaging allogenic responses of decidual T cells to EVT are prevented by immune suppressive mechanisms. The major decidual immune population, dNK, recognise and respond to EVTs using a range of receptors including maternal killer immunoglobulin-like receptors (KIR), which recognise the sole polymorphic HLA molecule expressed by EVTs: HLA-C^8^. Certain combinations of (KIR) and fetal HLAC variants are associated with pregnancy disorders such as preeclampsia, where trophoblast invasion is deficient^9^. However, the precise cellular interactions in the decidua supporting early pregnancy are poorly understood.

Due to the limited prior knowledge of this tissue in early pregnancy, we employed an unbiased approach to comprehensively resolve cell states involved in maternal–fetal communication at this interface. Furthermore, we aimed to decode cell-cell communication networks through systematic analysis of surface receptor-ligand analysis in a quantitative manner. Single–cell RNA sequencing (scRNA-seq) allows us to identify cell populations, and the many receptors and ligands expressed by these cells.

Molecular analysis of cell-cell communication requires a tool that takes into account the complexity of multi-subunit receptors and their ligands, and discerns the specificity of the interactions. Therefore, we combine single-cell transcriptomics with a computational framework to predict cell-type specific ligand-receptor complexes with a new database: www.CellPhoneDB.org.

By integrating these predictions with spatial *in situ* analysis, we generate a detailed molecular and cellular map of the human decidua and placenta, which provides new insights into the functional organisation of the fetal-maternal interface. Our findings reveal unexpected cellular heterogeneity and interactions between fetal trophoblast cells, maternal fibroblast cells and multiple dNK cell subsets, orchestrating the support of early pregnancy.

## Results

### Analysis strategy

Here, we combine high-throughput and in-depth transcriptome profiling of single cells to generate a comprehensive cellular map of the fetal-maternal interface. Cell suspension from human decidua and placenta and matched maternal peripheral blood from early pregnancy (6–14 weeks) were analysed by single cell RNA-sequencing (scRNA-seq) using droplet-based cell encapsulation (10x Genomics Chromium) as well as the plated-based Smart-seq2 (SS2) method for focused profiling^10^ (Fig. 1a). In order to identify relevant cell surface and secreted receptor-ligand pairs, we develop ‘CellPhoneDB’, a repository, available at www.CellPhoneDB.org (Fig. 1b) and an accompanying algorithm to predict cellular interactions.

**Fig. 1.**
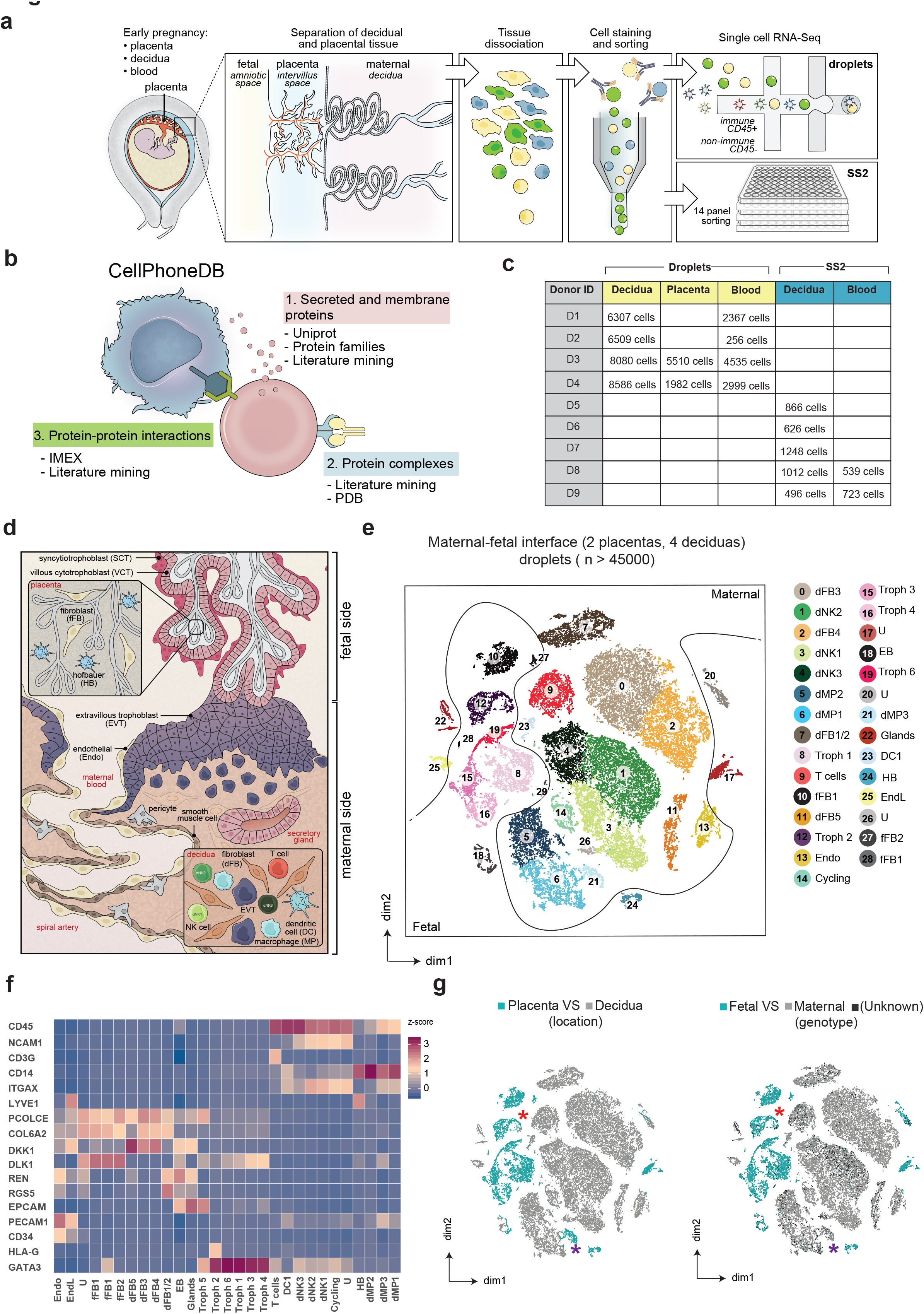
Identification of novel cell types at the maternal-fetal interface. **a**, Workflow of collection and processing of fresh, first trimester, decidual and placental tissues and maternal peripheral blood from first trimester elective terminations, for scRNA-seq analysis. B cells are not present in the decidua^56^ (Extended Data Fig. 4a) and were excluded from the analysis. **b**, Information stored in CellPhoneDB, www.CellPhoneDB.org (see Methods). **c**, Total numbers of cells that passed the quality control, processed by SS2 and droplet scRNA-seq. **d**, Diagram illustrating the anatomy of the placental-decidual interface in early pregnancy. **e**, Placental and decidual cell clusters obtained by the droplet-encapsulation method and visualised by tSNE. Colors indicate cell type/state, assigned by unbiased clustering and manual annotation (see Methods). Acronyms are listed in Fig. 1d. **f**, Heatmap showing the zscore of the mean log-transformed normalised counts per cluster of selected key genes used to identify general clusters in the different cells. For a more extensive set of genes see Supplementary Table 2–3 and Extended Data Fig. 2. **g**, Panels indicating the origin of cells in the tSNE plot shown in Fig. 1e. The left panel shows placental (blue) versus decidual (grey) cells and the right panel shows fetal (blue) versus maternal (grey) cells. Unassigned cells are colored in dark grey. The blue star highlights maternal cells from the placenta and the red star indicates decidual cells of fetal origin.

### Maternal and fetal cells present in decidua and placenta in early pregnancy

After computational quality control (see Methods) of cells profiled by droplet encapsulation, we obtained single-cell transcriptomes from four decidual samples (29,482 cells), matched peripheral blood mononuclear cells (PBMC) (10,157 cells), and two matched placental samples (7,492 cells) (Fig. 1c, Extended Data Fig. 1a). To generate a census of all the decidual and placental cells in the uterus, we performed unsupervised graph-based clustering of the combined dataset and used cluster-specific marker genes to annotate the clusters (Fig. 1d-f, Extended Data Fig. 1–2). Cells were confirmed as being fetal or maternal in origin by evaluating scRNA-seq reads overlapping single nucleotide polymorphisms (SNPs) called from maternal or fetal genomic DNA (Fig. 1g, Extended Data Fig. 1c-d). Decidua and placenta were mainly comprised of maternal and fetal cells respectively, however there were small numbers of fetal *HLA-G+* EVTs present in the decidua, and maternal myeloid cells (MP3 cluster) were detected in the placenta; these are likely to be patrolling maternal monocytes that attach to the syncytial surface^11^.

The decidua is composed of glands, vessels, stroma, and immune cells. Differentially expressed genes allowed us to identify subpopulations as follows (Fig. 1f, Extended Data Fig. 2). The cluster of decidual glandular epithelial cells is characterized by high levels of the epithelial marker *EPCAM* and the progestagen associated endometrial molecule (*PAEP*). We identify endothelial cells expressing *CD34* and *CD31* antigen (*PECAM1*). A heterogeneous population of fibroblasts express matrix protein genes (e.g. *PCOLCE, COL6A2*) and is negative for *PAEP, CD45*, and *CD31*. We reveal the presence of three dNK subsets (the most prevalent immune populations), two macrophage subsets, one dendritic cell subset (DC1). Decidual CD4+ and CD8+ T cells cluster together (Fig. 1e).

In the placenta, we identify trophoblast cells, which lack the expression of histocompatibility complex A (*HLA-A*) and B (*HLA-B*) and fetal fibroblasts expressing the notch ligand *DLK1*, known to be present in the core of the placental villi^11^ (Fig. 1f, Extended Data Fig. 2). We identify the placental myeloid cells, Hofbauer cells, which express lymphatic endothelial marker *LYVE-1* and the *CD14* molecule^12^.

### Placental trophoblast cells differentiate along two main pathways

Placental development is characterised by fetal trophoblast differentiation into EVTs and syncytiotrophoblast (SCT). The molecular transition and regulation of this process in humans is poorly characterised. We leveraged our scRNA-seq dataset to reconstruct the developmental relationship of all trophoblast cells present in both placental and decidual samples using the pseudotime trajectory algorithm Monocle 2^13^. In agreement with previous reports^14^, we resolved two distinct pathways for SCT and EVT differentiation (Fig. 2a).

**Fig. 2.**
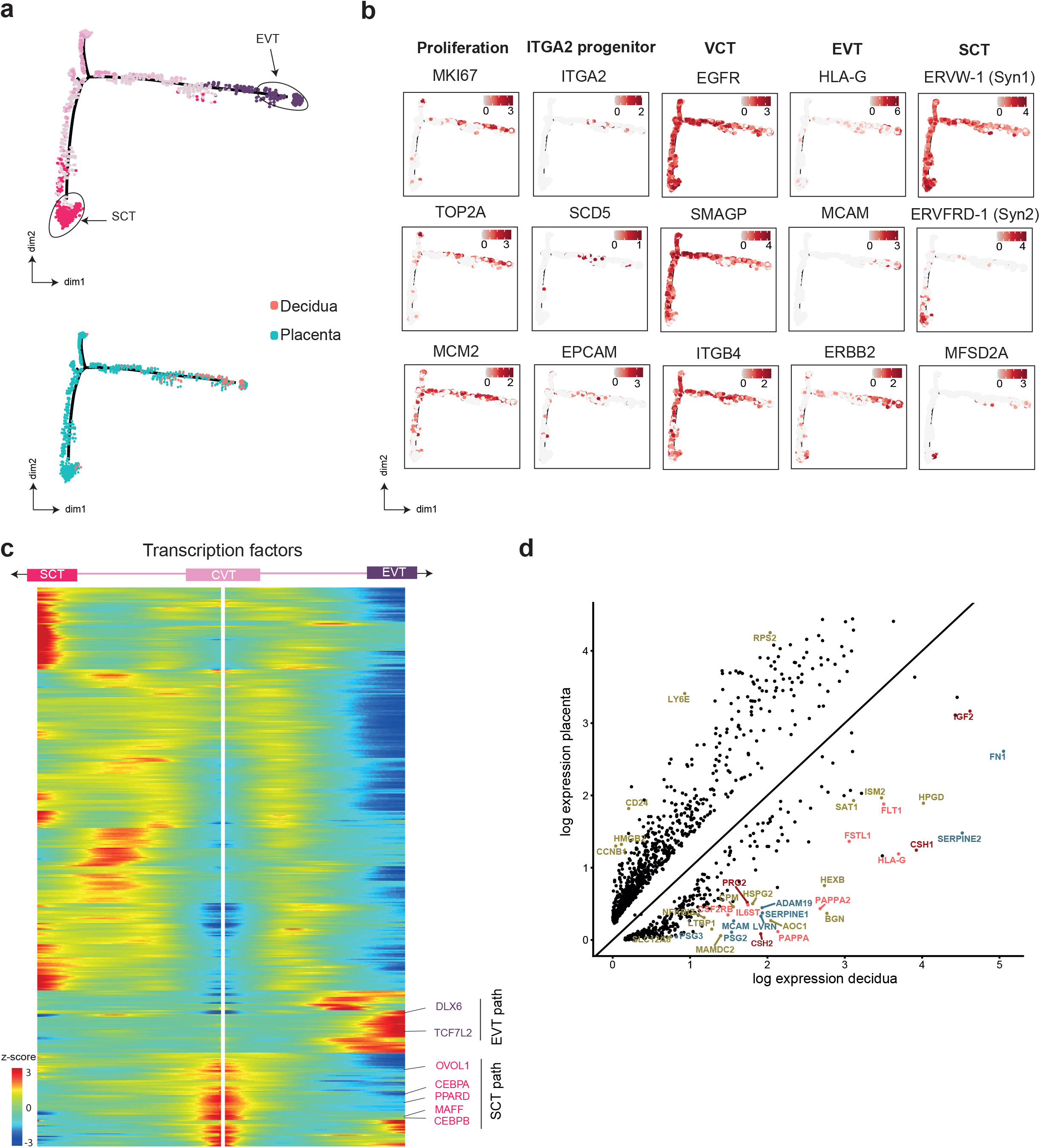
Differentiation of trophoblasts defined by pseudotime inference. **a**, Pseudotime ordering of cells reveals two distinct trophoblast differentiation pathways, SCT and EVT. Placental and decidual origin of cells is shown in the lower panel. **b**, Marker genes defining the two trophoblast differentiation pathways shown in Fig. 2a. Genes involved in the cell cycle are also displayed. **c**, Heatmap showing transcription factors identified as varying significantly along the trajectory (*q*-value<10^−4^) from the VCT (center) to EVTs (right) or SCT (left). Transcription factors relevant for EVTs and SCT are highlighted. **d**, Differential expression analysis between the decidual and placental EVT.

Villous cytotrophoblasts (VCTs) fuse to become SCT and this is mediated by expression of Syncytin-2 (*ERVFRD-1*) binding to its receptor MFSD2A. We observed a population of cells expressing *ERVFRD-1* prior to SCT fusion (Fig. 2b). Furthermore, *ERVFRD-1* expression declines in cells expressing the receptor *MFSD2A* at the end of the SCT differentiation trajectory, indicating some syncytial cells are present. *HLA-G*+ EVT are most abundant in placental samples with small numbers in decidua (Fig. 2a). In our placental isolates, approximately 40% of *HLA-G*+ EVT express the cell cycle gene *MCM2*; in contrast, none of the decidual EVT are cycling. Our analysis of trophoblast differentiation reveals the transcription factors (TFs) involved in the differentiation of EVTs and SCT fusion. The data confirms involvement of the TF *OVOL1^15^* and predicts many new factors involved in trophoblast fusion (Fig. 2c, Extended Data Fig. 3). Our pseudotime reconstruction also captures a proliferative trophoblast population (Fig. 2b), characterised by expression of *ITGA2, SCD5* and *EPCAM* as described previously^1^6.

Little is known about differentiation of EVTs as they migrate into the decidua. To investigate decidual EVTs further, we enriched for HLA-G+ CD45- EVT cells in the decidua and performed SS2 scRNA-seq from an additional five decidual samples (Extended Data Fig. 4). We then compared the transcriptomes of placental EVTs (in the cytotrophoblast cell columns) with decidual EVTs (interstitial trophoblast). This reveals a number of cell communication and adhesion molecules, as well as hormones characterising decidual EVTs (Fig. 2d). Other molecules that are upregulated as EVTs invade include: the *IGFBP* dependent molecule *PAPPA, PAPPA2*, and the tyrosine kinase *FLT1*. These are all potential biomarkers used to predict pregnancy complications^17,18^.

### Heterogeneity of stromal and glandular epithelial cells defines two decidual layers

We describe five clusters of decidual fibroblast cells (dFB1–5), expressing extracellular matrix molecules (e.g. *PCOLCE, COL6A2*) and lacking markers for endothelial cells (*CD34*), trophoblast (*HLA-G*) and epithelial cells (*EPCAM*) (Fig. 3a, Extended Data Fig. 5). dFB1 cells express the canonical pericyte markers *MCAM* (*CD146*) and *PDGFRB* (*CD140b*), as well as *RENIN* and *RGS5*, both involved in regulating vascular tone (Fig. 3b). The dFB2 population shares an expression pattern with dFB1, such as smooth muscle genes (e.g. *MYH11*), and the matrix protein *MGP*. dFB2 cells also have features of specialised smooth muscle cells involved in tissue remodelling, with higher levels of the matrix metalloprotease *MMP11* and the cathepsin *CTSK*.

**Fig. 3.**
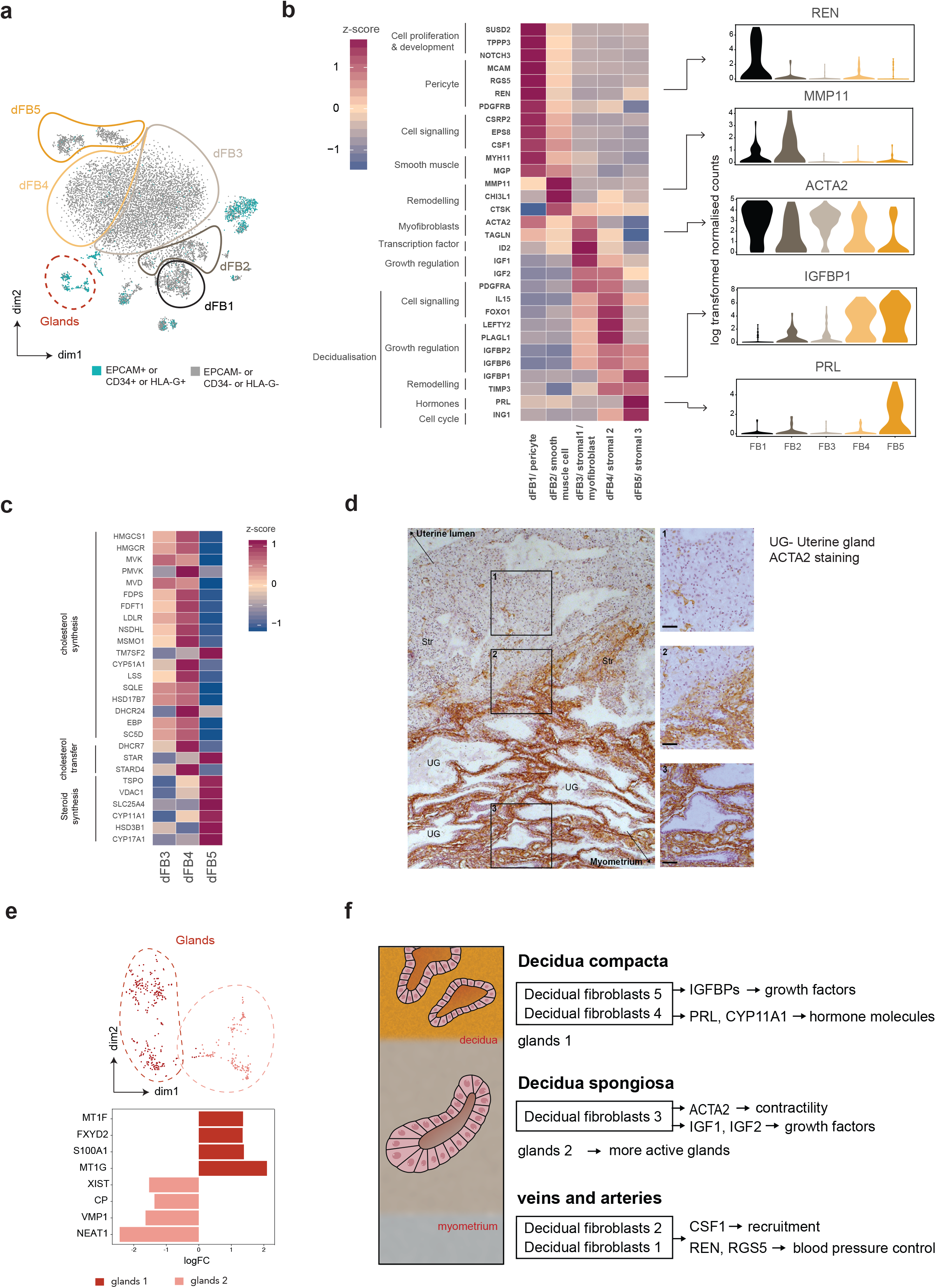
Single-cell transcriptomics and *in situ* staining separates the decidual mucosa into two distinct layers. **a**, tSNE plots showing cells with high expression levels (z-score > 2) for endothelial cells (*CD34*), EVT (*HLA-G*) and epithelium (*EPCAM*) cells that were not considered for the subsequent analysis of fibroblasts. **b**, Heatmap shows selected genes for the five dFBs subpopulations. The right panel shows log-transformed normalised counts for selected genes. **c**, Heatmap showing differential expression of enzymes involved in cholesterol and steroid synthesis in the three stromal cell subsets. **d**, IHC of decidual sections stained for ACTA2. ACTA2+ stromal cells are confined to the stromal cells of the deeper decidua spongiosa whilst those in the decidua compacta are ACTA2-. The scale bar represents 50 µm. **e**, (above) tSNE plots showing two clusters of glandular epithelial cells; (below) bar plots showing selected differentially expressed genes between these two clusters of epithelial glands. **f**, Diagram illustrating the structure of the two layers of the maternal decidua during pregnancy.

The dFB3–5 cell populations all express the WNT inhibitor *DKK1*, known to mark decidual stromal cells^19^. dFB3 shares myofibroblast features with dFB1/2 showing high levels of smooth muscle *ACTA2* and *TAGLN* with dFB1/2. Members of the insulin growth factor (IGF) family are differentially expressed in the dFB3–5 subsets, with dFB3 expressing higher levels of *IGF1/2*, and dFB4/5 expressing high levels of *IGFBP1/2/6*. In contrast, the dFB4/5 populations are decidualised stromal cells, expressing higher levels of the metallopeptidase inhibitor *TIMP3* and *IGFBP1* with dFB5 expressing high levels of the pregnancy hormone prolactin (*PRL*), used as a marker of decidualisation *in vitro*. dFB5 cells express the cell cycle inhibition protein *ING1*, further supporting this population as being the most differentiated. Pseudotime analysis predicts a differentiation trajectory from dFB1 to dFB5, positioning dFB5 at one end of the progression (Extended Data Fig. 6). This trajectory links the three dFB3–5 populations as expected. More surprisingly, it suggests possible interrelationships between and plasticity across all five dFB subpopulations. The pericyte-like population (dFB1), is positioned at the other end of the trajectory, confirming reports that pre-decidual stromal cells express pericyte markers cells^20^. Indeed, dFB1 cells seem able to self-renew as they express genes involved in cell proliferation and development (e.g. *TPPP3* and *NOTCH3*).

Steroid hormones play an important role in pregnancy^21–23^, however their presence cannot be directly measured by gene expression analysis, as they are organic compounds rather than proteins. Instead, we analysed expression of the steroid biosynthesis pathway genes. Enzymes involved in cholesterol synthesis are upregulated in dFB3/4, and downregulated in dFB5, which expresses enzymes involved in pregnenolone synthesis (Fig. 3c). This suggests that, during differentiation from dFB3/4 to dFB5, a metabolic transition occurs, such that dFB5 cells synthesize both steroids and prolactin.

Pericytes and smooth muscle cells (dFB1/2) encircle vessels, whilst stromal cells (dFB3–5) are located throughout the decidua, in close contact with maternal immune cells and invading EVTs. The gradual increase of decidualisation molecules in dFB4/5 suggest a microanatomical zonation, and a distinct distribution of dFB3 and dFB4/5 within the decidua. Histologically, the decidua is divided into two morphological zones: decidua spongiosa close to the myometrium, and the more superficial decidua compacta. In order to locate the stromal subsets (dFB3–5), we stained tissue sections for the marker ACTA2, expressed by dFB3 and downregulated in dFB4/5 (Fig. 3d). ACTA2 expression clearly divides the decidua stroma into two defined layers corresponding to the decidua spongiosa (dFB3) and compacta (dFB 4/5).

The EPCAM+ epithelium of uterine glands present is also histologically distinguishable in the two decidual layers. In the decidua spongiosa glandular cells are ‘plump’, with abundant cytoplasmic content, described as hypersecretory^24^. In contrast, in decidua compacta the glandular epithelium forms a thin layer, and appears ‘exhausted’. Our single-cell transcriptomics analysis resolves two different glandular epithelial cell clusters; one expressing transporters (e.g. *FXYD2, MT1G*), suggestive of more active metabolism and high secretory function of spongiosa cells, and the other with high expression of mitochondrial genes and molecules involved in autophagy (e.g. *TMEM49*) (Fig. 3e).

By integrating scRNA-seq with histology, we assign the location of the stromal and glandular epithelial cells to specific microanatomical zones within human decidua (Fig. 3f).

### Immune cell profile in the decidua during early pregnancy

To further define the heterogeneity of maternal decidual immune cells and evaluate TCR and KIR receptor sequences, we generated a full-length transcript scRNA-seq dataset using our robotic SS2 platform. We profiled 4,248 CD45+ cells in decidua and 1,262 cells from maternal blood, isolated using antibodies against proteins characteristic of decidual immune cells^25–27^ (Fig. 4a, Extended Data Fig. 4). We integrated these data with our droplet-based dataset by applying a computational approach for scRNA-seq alignment across protocols^28^ (Fig. 4a-b), and assigned cluster identities based on the most highly differentially expressed genes for each cluster. The clusters were separated into cells resident in decidua versus populations of leukocytes present in both blood and decidual samples (Extended Data Fig. 7).

**Fig. 4.**
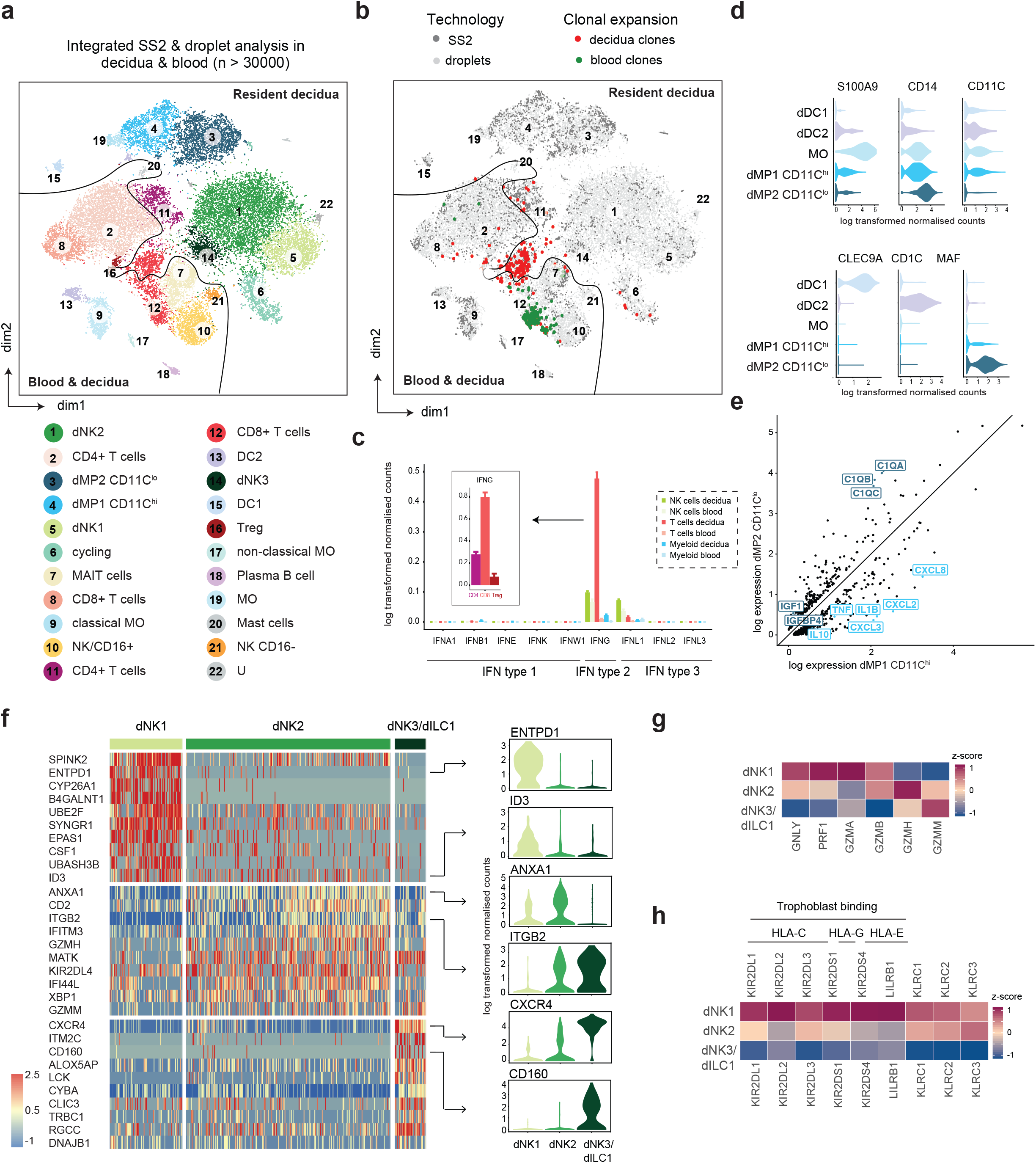
Single-cell transcriptomics defines several populations of decidual immune cells. **a**, tSNE plot showing matched maternal decidual and peripheral blood immune cells. Colors indicate cell type/state, assigned by unbiased clustering and manual annotation (see Methods). **b**, tSNE plot showing the integration of SS2 and droplet RNA-seq data, and T cells with expanded TCR clonotypes. **c**, Mean log-transformed normalised expression of members of the IFN pathway in the maternal immune subsets. Error bars represent standard error of the mean. **d**, Violin plots showing log-transformed normalised expression of several myeloid markers. **e**, Differential expression analysis between the two main mononuclear phagocyte populations (*q*-value<0.01). **f**, (Left) Heatmap showing z-scores of the log-transformed normalised expression of markers defining the three dNK subsets. (Right) Violin plots of markers of dNK1–3 subsets. **g**, Heatmap showing z-scores of the mean log-transformed normalised expression of markers of lytic secretory granules, as expressed on dNK subpopulations. **h**, The z-scores of KIR receptors mean expression levels. Expression values were generated using KIRid approach (see Methods).

Compared to blood, CD4+ and CD8+ T cells population are present at lower frequencies in the decidua. Decidual T cells have features of tissue-resident T cells, such as expression of *CD69, CXCR3* and *CXCR4* (but are negative for *SELL* and *CCR7*) (Extended Data Fig. 7, 8a). CD4+ T cells have attributes of a Th1 phenotype (*IFNG*+, *PTGDR2-*), although the sparse expression of *RORC, KIT* and *IL17* indicates the presence of a few ILC3 and Th17 cells in this cluster (Extended Data Fig. 7, 8a). A small subset of FOXP3+ Tregs from both decidua and blood cluster together.

Decidual CD8+ T cells share similarities with a specific blood memory CD8+ T cell subset^29^, which express high levels of genes encoding granzyme A and B, perforin and granulysin, and lack expression of *IL7R*. Both populations are clonally expanded, with identical clonotypes found in both decidua and blood (Fig. 4b, Extended Data Fig. 9). In contrast to blood T cells, decidual T cells (and not NK cells) have the highest expression of Type II IFN (*IFNG*), and our CellPhoneDB approach shows that the corresponding receptor (*IFNR*) is widely expressed in different subsets of immune and non-immune cells (Fig. 4c, Extended Data Fig. 8b).

The myeloid compartment in the decidua is composed of two macrophage populations characterized by high expression levels of *CD14* and *MHCII* genes. These two populations are distinct in terms of their expression of *CD11C*, termed *CD11C*^hi^ dMP1 and *CD11C*^lo^ dMP2. We identify two populations of dendritic cells (DC) expressing classical markers for DC1 and DC2 (Fig. 4d). Mast cells are a minor population (∼0.5% of total immune cells in the decidua).

The dMP1 *CD11C*^hi^ population shares genes with decidual monocytes (e.g., *S100A9*), and expresses macrophage markers such as *CD163, CD68* and *CSF1R*, suggesting a monocyte-derived macrophage identity. In addition, expression of inflammatory (*IL1B, TNF, IL6, CXCL2,3,8*) and anti-inflammatory cytokines (*IL10*) by this dMP1 population highlights the balanced expression of genes with a role in remodelling and tolerance induction (Fig. 4e).

In contrast, *CD11C*^lo^ dMP2 cells express higher levels of macrophage markers, the mannose receptor *MRC1* (*CD206*), *MAF* and *F13A1*, consistent with a more differentiated tissue-resident macrophage phenotype, as previously described in human skin^30^. Secretion of complement-related factors (*C3, C1QA, CD1QB, C1QC*) and growth factors (*IGF1, IGFBP4*) by dMP2 suggest a role in tissue clearance and growth. Some of the dMP2 cells are cycling (*i.e*. expressing *MKI67*), indicating their potential to self-renew locally (Extended Data Fig. 7).

### Three main decidual natural killer (dNK) subsets

The dominant population of immune cells in the first trimester decidua are CD56+ NK cells, expressing tissue resident markers *CD49a* (*ITGA1*) and *CD9* (Extended Data Fig. 8a). We computationally identify three main subsets of decidual NK cells (dNK1–3), which are all distinct from peripheral blood NK cells (Fig. 4a). The markers that best discriminate between dNK1–3 clusters are: the surface protein CD39 (ENTPD1) and the transcription factor *ID3* for dNK1 cells; the annexin member *ANXA1* and the integrin *ITGB2* for dNK2 cells; and chemokine receptor *CXCR4* and the glycoprotein *CD160* for dNK3 cells (Fig. 4f). It is noteworthy that the dNK3 population exhibits features of ILC1-like cells, so we call them dNK3/dILC1. The typical ILC1 markers that are expressed in these cells are *TBX21, CD160, CD161* (*KLRB1*) and *CD103* (*ITGAE*), but they are negative for *CD127* (*IL7R*) (Extended Data Fig. 8a).

The two dNK populations, dNK1 and dNK2, differ in the content of their secretory granules, only expressed at low levels in dNK3/dILC1 cells. dNK1 express high levels of cytotoxic granule molecules (*GNLY, GZMA, GZMB*), whilst dNK2 and dNK3/dILC1 are characterised by the unconventional granzymes *GZMH* and *GZMM* (Fig. 4g). It is interesting to note that dNK1 cells express *CYP26A1*, known to alter the composition of murine dNK subsets^31^ (Fig. 4f).

Receptors of the Killer-cell immunoglobulin-like Receptor (KIR) family that bind to HLA class I molecules, and other NKR binding to EVT ligand, are differentially expressed on dNK1 and dNK2 cells. KIR recognise polymorphic HLA-C allotypes, and, because KIR family members are also polymporphic and highly homologous, quantifying expression of individual KIR genes is challenging^9^. To solve this problem, we used our full-length transcript SS2 data to map each donor’s single cell reads to the corresponding donor-specific reference of KIR alleles, based on existing haplotypes (see Methods). The levels of expression of the specific KIR receptors that interact with HLA-C (*KIR2DL1/DL2/DL3* and *KIR2DS1/DS4*), and the HLA-G- binding receptor *LILRB1*, are high on dNK1, low on dNK2, and absent on dNK3/dILC1 (Fig. 4h). Both dNK1 and dNK2 have high levels of the HLA-E interacting receptors (*KLRC1, KLRC2, KLRC3*), which include both activating and inhibitory receptors (inhibitory KLRC1, activating KLRC2 and KLRC3).

In summary, our transcriptomic analysis reveals three subsets of dNK cells with potentially different regulatory, inhibitory and cytotoxic capacity.

### dNK subsets have distinct receptor-ligand interactions

Many of the decidual cell populations described above express specific cell surface receptor and ligand molecules, which we curate in CellPhoneDB. We take into account the expression levels of ligands and receptors within each cell type, and calculate statistically significant ligand–receptor pairs by empirical shuffling (Fig. 5a). This way, we predict functional interactions between cell populations *via* specific protein complexes. Here we focus on receptors and ligands that are differentially expressed by dNK subsets to predict their significant cellular interactions (Fig. 5b).

**Fig. 5.**
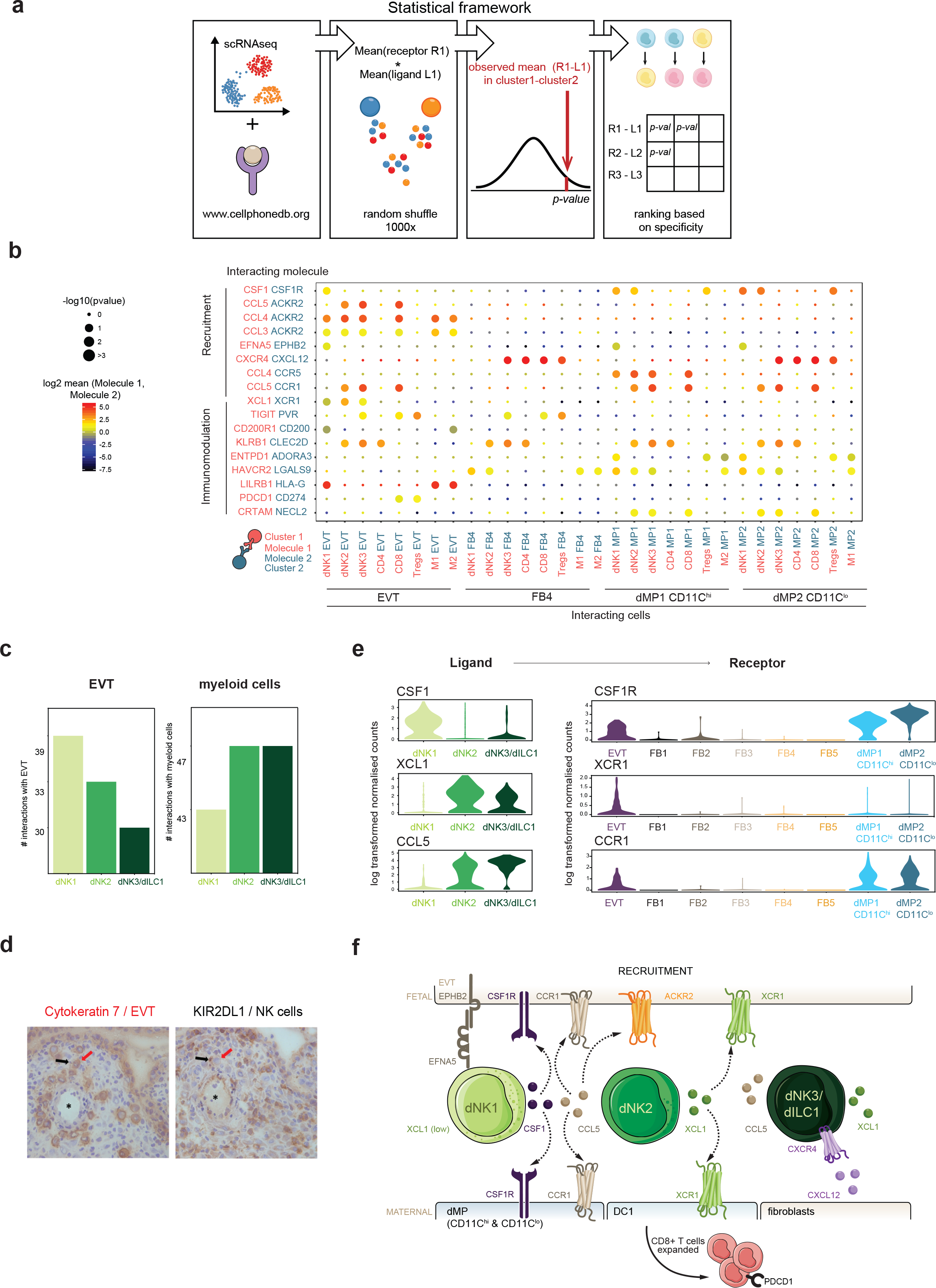
Cell-cell interactions at the maternal-fetal interface inferred from CellPhoneDB. **a**, A statistical framework for inferring ligand/receptor pairs specific to two cell types from single cell transcriptomics data. The *p*-values for predicted receptor-ligand pairs across two cell type clusters are calculated using permutations, where cells are randomly re-assigned to clusters (see Methods). **b**, Overview of selected receptor/ligand interactions, with *p*-values indicated by the size of the circles. The means of the average receptor expression levels of a cluster and its interacting cluster are indicated by colour. **c**, Total numbers of the interactions between the three decidual NK subsets, EVT and myeloid cells. **d**, IHC images of decidual serial sections of the decidual implantation site (10 weeks gestation), stained for the trophoblast cell marker, Cytokeratin-7 (red arrow) and the inhibitory NK receptor, KIR2DL1 (black arrow). The asterisk (*) marks the lumen of a spiral artery that supplies the conceptus. **e**, Violin plots showing log-transformed normalized expression level of several ligands and receptors pairs inferred by our statistical framework. **f**, Diagram of the main receptors and ligands expressed on the three dNK subsets that are involved in cellular recruitment.

Our analysis reveals more potential interactions for dNK1 with EVTs than for the two other dNK populations (39 dNK1 vs 33 dNK2 and 30 dNK3/ILC1). This suggests that EVTs preferentially bind the dNK1 subset (Fig. 5c). Specific interactions include *LILRB1* - *HLA-G* and *KIRs* - *HLA-C*. The physical proximity between dNK1 and EVT in the decidua basalis is confirmed by IHC of serial sections stained for CK7 (EVT) and KIR2DL1 (dNK1) (Fig. 5d). In addition, our method predicted different growth factor and chemokine interactions for dNK1/2 and dNK3/dILC1 cells. dNK1 expresses higher levels of the cytokine *CSF1*, whose receptor, *CSF1R*, is expressed by EVTs and macrophages (Fig. 5e). Secretion of CSF1 by dNK cells, and the impact of their interaction with the CSF1R on EVT has been described^32,33^. We are now able to pinpoint that this interaction is mediated specifically by the dNK1 subset.

In contrast, dNK2–3 express low levels of *CSF1*, and high levels of *XCL1* and *CCL5*. EVT strongly express the chemokine-scavenging receptor *ACKR2^34^*, that binds numerous inflammatory CC-chemokines, including *CCL3, 4* and *5* (Fig. 5e). ACKR2 acts as a decoy receptor, effectively putting a brake on the inflammatory response^35,36^. *CCL5* also binds the chemokine receptor *CCR1* expressed by EVT, and may regulate EVT migration into the decidua^37^. *XCL1* binds the *XCR1* receptor expressed by both EVT and decidual myeloid cells.

### Multiple anti-inflammatory, tolerogenic interactions at the maternal-fetal interface

The diversity and large number of receptor-ligand interactions between EVT and decidual cells must relate to the development of an anti-inflammatory, tolerogenic environment, critical for the peaceful compromise needed to define the territorial boundary between mother and fetus. We use CellPhoneDB to predict tolerogenic crosstalk in the maternal–fetal interface and identify the following statistically significant interactions. Interactions between EVT and dNK1 include the *CD200R1* on dNK1, potentially interacting with *CD200* expressed by EVT (Fig. 5b, 6a). EVTs also express inhibitory receptors, *CLEC2D* and *PVR*, whose ligands *KLRB1* and *TIGIT* are highly expressed on dNK2 and dNK3 respectively, providing additional NK-EVT interactions that could inhibit NK killing of EVTs.

**Fig. 6.**
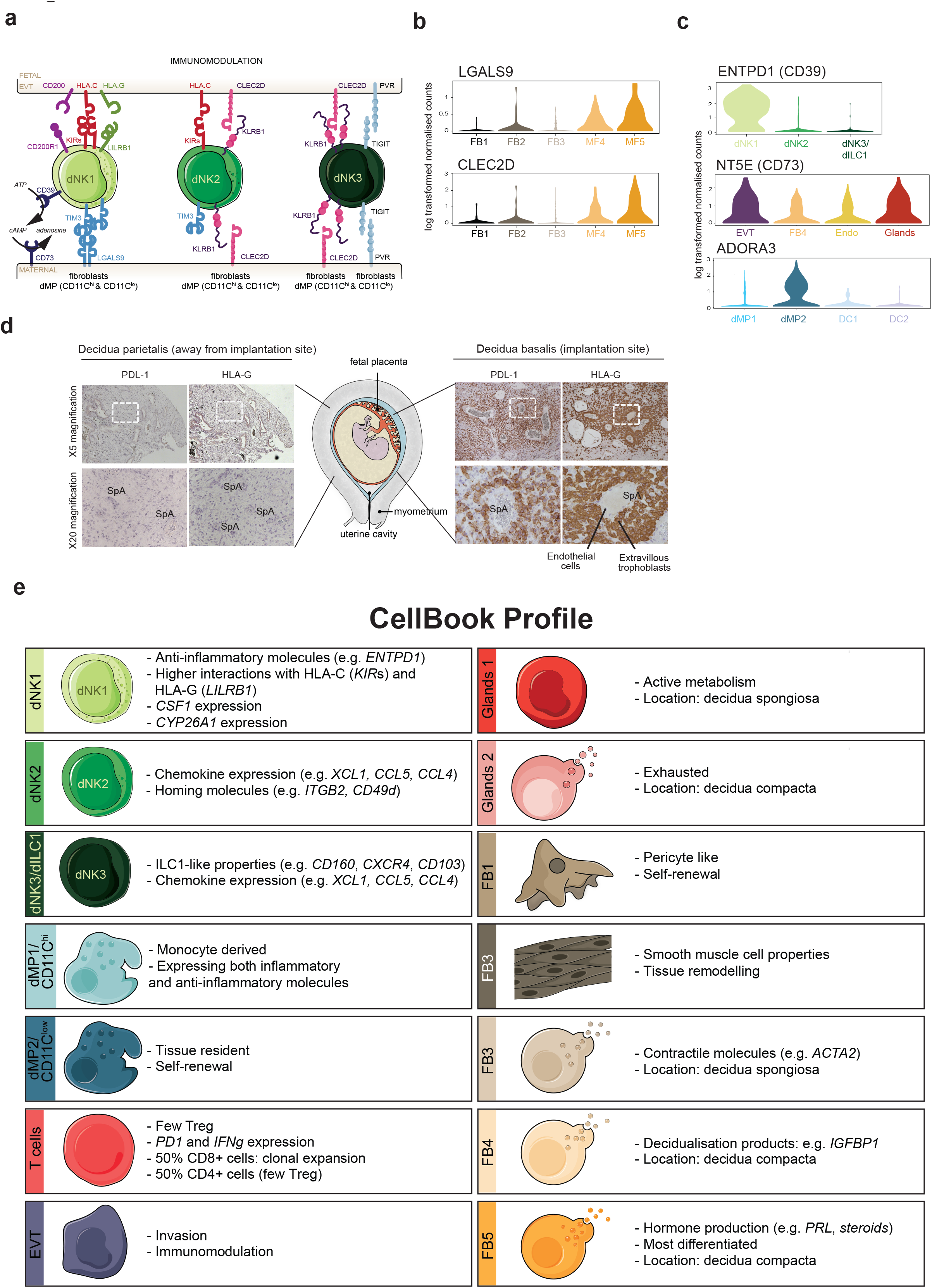
Multiple interactions dampen immune and inflammatory responses at the site of placentation. **a**, Interactions of the main receptors and ligands expressed on the three dNK subsets with their ligands on EVT and other maternal decidual cells that are involved in immunomodulation. **b**, Violin plots showing log-transformed normalized expression of several receptors in the decidual fibroblast populations. **c**, Violin plots showing logtransformed normalized expression of *CD39/CD73/ADORA3* in several cells in the decidua including EVT. **d**, IHC images of two decidual sections stained for the EVT marker, HLA-G and the inhibitory ligand PDL1. Lower images are areas in white boxes at higher power. HLA-G+ cells are only present at the site of placentation (decidua basalis) and are absent elsewhere (decidua parietalis). SpA identifies spiral arteries. The EVT is strongly PDL1+. **e**, Diagram showing main cell types in the decidua found in our analysis.

We predict that EVT cells interact specifically to suppress T cell responses. As mentioned above, the clonal expansion of decidual CD8+ T cells suggests they are responding to local APCs, and CD8+ T cells express the gene *PD1*. We note the high expression of *PDL1* in the EVT, which we confirm by IHC on serial sections stained for PDL1 and HLA-G (Fig. 6d). No other cells in the decidua are PDL1+, as shown by our IHC analysis of decidua parietalis away from the implantation site.

We hypothesize that the three differentiated stromal cell populations dFB3–5 also play an important role in contributing to the anti-inflammatory environment of the decidua (Fig. 6c). In the superficial decidua compacta where EVTs first invade, dFB4/5 cells express high levels of Galectin-9 (*LGALS9*), with the cognate receptor *TIM3* expressed on dCD8+ T cells and dNK1/2 cells. Like EVT cells, dFB4/5 cells express high levels of the c-type lectin (*CLEC2D*), whose ligand *KLRB1* is expressed on dNK2 and dNK3/dILC1 cells (Fig. 5b). The steroid biosynthesis in dFB5 cells may also contribute to the tolerogenic environment in the decidua compacta.

In addition to EVT and stromal cells, immune subsets themselves also contribute to the tolerogenic environment. For instance, dNK1 cells express *SPINK2*, an inhibitor of inflammatory and secreted kallikrein molecules^38^, and dNK2/3 express high levels of the annexin member *ANXA1*, with anti-inflammatory roles. The dNK1 subset also express *ENTPD1* (*CD39*), which together with *CD73* (*NT5E*) converts ATP to adenosine, a small molecule known to abrogate immune activation^39^. The expression of *CD73* is high in epithelial glands and EVTs, and the adenosine receptor (*ADORA3*) is found on decidual myeloid cells (Fig. 5b, 6c).

In summary, both EVT and stromal cells, as well as innate immune cell populations, all contribute to setting up an anti-inflammatory environment at the site of placentation in early pregnancy. This is mediated by multiple classes of signalling molecules: cell surface tethered receptor-ligand complexes, secreted proteins, as well as small molecules (adenosine and steroid hormones).

## Discussion

Healthy pregnancy depends on the normal growth and development of the placenta in the first trimester, yet the cellular and molecular events occurring in the uterus are still not understood. Difficulties in accessing material and the absence of reliable *in vitro* models^40^, as well as substantial differences between human and rodent models, have hindered progress. Here, we overcome some of these limitations by profiling human placental and decidual tissue from early pregnancy (6–14 weeks) using a combination of scRNA-seq methods. By integrating this data with maternal and fetal whole genome sequencing, we can assign the fetal and maternal genotype of each individual cell. Thus, we are able to evaluate the contribution of maternal and fetal cells within both the placenta and decidua, and generate an unbiased map of this important tissue interface highlighting new cell states (Fig. 6e).

Trophoblast cells, derived from trophectoderm, are extra-embryonic cells providing a physical barrier between mother and fetus. In contrast to other studies that have focused on term placental tissue^14,41^, we have analysed trophoblast cells in early pregnancy at the time the placenta expands and becomes established. We were, for the first time, able to resolve trophoblast development into a tri-partite branching trajectory using first trimester samples, and confirmed that in humans there is differentiation along two main lineages, EVT and SCT. Genes known to be involved in fusion of VCT into the overlying syncytial layer are highly expressed at the end of the SCT trajectory. Our differential expression analysis along the trajectory allowed us to identify novel TFs involved in trophoblast differentiation and fusion. By comparing the HLA-G+ EVT cells from the placental villi with HLA-G+ cells isolated from decidual tissue, we highlight genes that are upregulated during the invasive process. These include genes such as *PAPPA* and *FLT1*; low *PAPPA* levels in maternal serum are associated with pregnancy disorders caused by poor trophoblast invasion and defective placentation^16,17^.

The combination of single-cell transcriptomics and spatial *in situ* methods provide a unique framework to study the microanatomical distribution of cells in the tissue^42,43^. We resolved five decidual fibroblasts subsets (dFB1–5) and two epithelial glands that we located in the two layers of the decidua, spongiosa and compacta. Two populations, dFB1 and dFB2, have features of the pericytes/medial smooth muscle cells that surround the crucial vessels supplying the conceptus and regulating vascular tone, as suggested by previous work^20^. dFB3–5 are fibroblasts with characteristics of decidual stromal cells, with the most differentiated hormone-secreting cells (dFB5) present in the superficial decidua compacta, whilst the ACTA2+ population (dFB3) surrounds the hypersecretory glands of the underlying decidua spongiosa. Their potential contractile properties may help deliver the glandular products into the villous placenta to provide nutrition before hemochorial nutrition is established at around 8–10 weeks’ gestation.

During early pregnancy, the decidua displays an unusual composition of maternal immune cells, dominated by dNK cells that we can now separate into three distinct populations. Our study highlights clonally expanded CD8+ T cells, an equal proportion of CD4+ T cells, and a smaller population of Tregs. We find a range of myeloid cells including macrophages and classical DCs, and mast cells and B cells are virtually absent. The dNK3 resemble ILC1^44–46^, although they do have decidual-specific features, such as a lack of *CD7* and *IL7R* (*CD127*) expression. dNK1 are of particular interest since they express NK receptors that bind to HLA-C, HLA-G and HLA-E expressed by EVT^40^. We show high expression of cytotoxic molecules of secretory granules for dNK1, although the lack of expression of b2 integrin (*ITGB2*) would reduce their cytotoxic potential^47^. dNK1 cells express high levels of the cytokine *CSF1*, whose receptor is involved in trophoblast recruitment and activity^32,33^.

Our cell–cell communication analysis framework indicates that dNK1 cells have a unique regulatory profile. dNK1 cells express the ectonucleotidase *CD39* (*ENTPD1*) which degrades extracellular ATP (eATP) in tandem with *CD73* (*NT5E*), expressed in the non-immune decidual populations, to promote the synthesis of adenosine that suppresses both innate and adaptive immune responses^48,49^. dNK2 and dNK3 also express anti-inflammatory molecules like *ANXA1*, playing key immunoregulatory roles to protect the conceptus and ensure its survival. In addition, dNK2–3 express high levels of the cytokines *CCL5* and *XCL1*, with their receptors present on trophoblast cells, myeloid and DC1 cells. Binding of *XCL1* to *XCR1* has been implicated in the recruitment of DC1s in other contexts^50^. This is consistent with our observation of expanded DC1 compared to DC2 in the decidua, in contrast to other tissues where DC2 vastly outnumber DC1^51,52^.

In summary, we present a single cell resolution atlas of the human maternal fetal interface, describing a range of novel cell subsets, their developmental trajectories, and surface receptors potentially mediating critical interactions between cell populations. The methods and database we have created, publicly available at www.CellPhoneDB.org, will be widely applicable to study cell–cell communication in other systems. This study identifies many novel molecular and cellular mechanisms that are operating at the fetal–maternal interface, and these insights may improve the diagnosis and treatment of infertility and complications of pregnancy.

## Methods

### Patient samples

All tissue samples used for this study were obtained with written informed consent from all participants in accordance with the guidelines in The Declaration of Helsinki 2000 from multiple centres.

Human embryo, fetal and decidual samples were obtained from the MRC/Wellcome-Trust funded Human Developmental Biology Resource (HDBR^57^, http://www.hdbr.org) with appropriate maternal written consent and approval from the Newcastle and North Tyneside NHS Health Authority Joint Ethics Committee (08/H0906/21+5). HDBR is regulated by the UK Human Tissue Authority (HTA; www.hta.gov.uk) and operates in accordance with the relevant HTA Codes of Practice.

Peripheral blood from woman undergoing elective terminations were under appropriate maternal written consent and approvals from the Newcastle Academic Health Partners (reference NAHPB-093) and HRA NHS Research Ethics committee North-East-Newcastle North Tyneside 1 (Rec reference 12/NE/0395)

Placental and decidual tissue for immunohistochemistry were obtained from elective terminations of normal pregnancies at Addenbrooke’s Hospital between 6 and 12 weeks’ gestation, under ethical approval from the Cambridge Local Research Ethics Committee (04/Q0108/23).

### Isolation of decidual, placental and blood cells

Decidual and placental tissue were washed in HAMS F12 medium, macroscopically separated and then washed for at least 10 mins in RPMI or HAMS F12 medium respectively before processing.

Decidual tissues were chopped using scalpels into approximately 0.2 mm^3^ cubes and enzymatically digested in 15ml 0.4mg/mL collagenase V (Sigma, C-9263) solution in RPMI 1640 medium (ThermoFisher Scientific, 21875–034)/10% FCS (Biosfera, FB-1001) at 37°C for 45 min. The supernatant was diluted with medium and passed through 100um cell sieve (Corning, 431752) and then 40um cell sieve (Corning, 431750). The flow-through was centrifuged and resuspended in 5ml of red blood cell lysis buffer (Invitrogen, 00–4300) for 10min.

Each first trimester placenta was placed in a petri dish and the placental villi were scraped from the chorionic membrane using a scalpel. The stripped membrane was discarded and the resultant villous tissue was enzymatically digested in 70 ml 0.2% trypsin 250 (Pan Biotech P10–025100P)/0.02% EDTA (Sigma E9884) in PBS with stirring at 37°c for 9 min. The disaggregated cell suspension was passed through sterile muslin gauze (Winware food grade) and washed through with Hams F12 medium (Biosera SM-H0096) containing 20% FBS (Biosera FB-1001). Cells were pelleted from the filtrate by centrifugation and re-suspended in Hams F12. The undigested, gelatinous tissue remnant was retrieved from the gauze and further digested with 10–15 ml collagenase V at 1.0mg/ml (Sigma C9263) in Hams F12 medium/10% FBS with gentle shaking at 37°C for 10 min. The disaggregated cell suspension from collagenase digestion was passed through sterile muslin gauze and the cells pelleted from the filtrate as before. Cells obtained from both enzyme digests were pooled together and passed through 100um cell sieve (Corning, 431752) and washed in Hams F12. The flow-through was centrifuged and resuspended in 5ml of red blood cell lysis buffer (Invitrogen, 00–4300) for 10min.

Blood samples were carefully layered onto a Ficoll-paque gradient (Amersham, Buckinghamshire, UK) and centrifuged at 2,000 rpm for 30 min without brakes. Peripheral blood mononuclear cells (PBMCs), from the interface between the plasma and the Ficoll– Paque gradient, were collected and washed in ice-cold phosphate-buffered saline (PBS), followed by centrifugation at 2,000 rpm for 5 min. The pellet was resuspended in 5ml of red blood cell lysis buffer (Invitrogen, 00–4300) for 10min.

### Flow cytometry staining, cell sorting and single-cell RNA sequencing

Decidual and blood cells were incubated at 4°C with 2.5ul of antibodies in 1% FBS in DPBS without Calcium and Magnesium (ThermoFisher Scientific, 14190136). DAPI was used for live/dead discrimination. We used an antibody panel designed to enrich for certain population for single-cell sorting and single-cell RNA sequencing (scRNA-seq). Cells were sorted using a Becton Dickinson (BD) FACS Aria Fusion with 5 excitation lasers (355nm, 405nm, 488nm, 561nm and 635nm Red), and 18 fluorescent detectors plus forward and side scatter. The sorter was controlled using BD FACS DIVA software (version 7).

For single-cell RNA-seq using the plate-based SS2 protocol, we created overlapping gates that comprehensively and evenly sampled all immune cell population in the decidua (Extended Data Fig 4). B cells (CD19+ or CD20+) were excluded from our analysis, due to their absence in decidua^56^. Single cells were sorted into 96-well full-skirted Eppendorf plated chilled to 4C, prepared with lysis buffer consisting of 10 ul of TCL buffer (Qiagen) supplemented with 1% b-mercaptoethanol. Single-cell lysates were sealed, vortexed, spun down at 300g at 4°C for 1 min, immediately placed on dry ice, and transferred for storage at –80°C. The SS2 protocol was performed on single-cells as described previously^58,59^, with some modifications^52^. Libraries were sequenced aiming at an average depth of 1 million reads/cell, on an Illumina HiSeq 2000 with v4 chemistry (paired-end 75-bp reads).

For the droplet scRNA-seq methods, blood and decidual cells were sorted into immune (CD45+) and non-immune (CD45-) fractions. Only viable cells were considered, and B cells (CD19+/CD20+) were excluded from our analysis, due to its absence in decidua^56^. Placental cells were stained for DAPI and only viable cells were sorted. Cells were sorted into an Eppendorf tube containing PBS with 0.04% BSA. Cells were immediately counted using a Neubauer hemocytometer and loaded in the 10x-Genomics Chromium. 10x-Genomics v2 libraries were prepared according to manufacturer’s instruction. Libraries were sequenced aiming at a minimum coverage 50,000 raw reads per cell on an Illumina HiSeq 4000 (paired-end, Read 1: 26 cycles; i7 index:8 cycles, i5 index: 0 cycles. Read 2: 98 cycles).

### Immunohistochemistry (IHC)

Tissue sections of 4um were cut from formalin-fixed paraffin wax-embedded human decidual and placental tissues. Sections were dewaxed with Histoclear, cleared in 100% ethanol and rehydrated through gradients of ethanol to PBS. Sections were blocked with 2% serum (of species in which the secondary antibody was made) in PBS, incubated with primary antibody for overnight at 4 RT°C and slides washed in PBS. Biotinylated horse anti-mouse or goat anti-rabbit secondary antibody were used, followed by vectastain ABC-HRP reagent (Vector, PK-6100) and developed with di-aminobenzidine (DAB) substrate (Sigma, D4168). Sections were counterstained with Carazzi’s haematoxylin and mounted in glycerol/gelatin mounting medium (Sigma, GG1–10). Primary antibody was replaced with equivalent concentration of mouse or rabbit IgG for negative controls. Tissue sections were imaged using a Zeiss Axiovert Z1 microscope and Axiovision imaging software SE64 V4.8.

### Whole genome sequencing

Tissue DNA and RNA were extracted from fresh frozen samples using the AllPrep DNA/RNA/miRNA kit (Qiagen) following the manufacturer’s instructions. Libraries for whole-genome sequencing were constructed according to Illumina library protocols, and 100-base paired-end sequencing was performed on HiSeq X genome analysers to an average of 30× coverage.

### Single cell RNA-seq data analysis

Droplet-based sequencing data was aligned and quantified using the Cell Ranger Single-Cell Software Suite (version 2.0, 10x Genomics Inc)^61^ against the GRCh38 human reference genome provided by Cell Ranger. Cells with fewer than 500 detected genes and for which the total mitochondrial expression exceeded 20% were removed. Genes that were expressed in fewer than 3 cells were also removed. SmartSeq2 sequencing data was aligned with HISAT2^62^, using the same genome reference and annotation as the 10x data. Gene-specific read counts were calculated using HTSeq-count^63^. Cells with fewer than 1,000 detected genes and more than 20% mitochondrial content were removed. Further, genes expressed in fewer than 3 cells were also removed.

Downstream analysis such as normalisation, k-nearest neighbour graph clustering, differential expression analysis and visualisation, were performed using the R package Seurat^64^. tSNE analysis was performed using a perplexity of 30. Clusters were annotated using canonical cell type markers. We further removed cells we did not gate for (most likely maternal blood B cells and fetal brain tissue), clusters for which the top markers were genes associated with dissociation-induced effects^65^, or mitochondrial genes, and a fibroblast cluster with high expression of hemoglobin genes due to background contamination of cell free RNA. Droplet-based and SmartSeq2 data were integrated using the Seurat alignment workflow^28^. During the alignment procedure we included sample identity as a covariate in the *ScaleData* function of Seurat for the 10x CD45- cells.

Trajectory modelling and pseudotemporal ordering of cells was performed with the Monocle 2 R package^66^ (version 2.4.0). The most highly variable genes were used for ordering the cells in the trophoblast cells and the differentially expressed genes across the fibroblast clusters for the fibroblasts pseudotime analysis. To account for the cell cycle heterogeneity in the trophoblast subpopulations, we performed hierarchical clustering of the highly variable genes and removed the set of genes clustering with known cell cycle genes such as CDK1. For similar reasons, we removed the mitochondrial and ribosomal genes for the trajectory analysis of the fibroblasts cells.

### Inferring maternal/fetal origin of single cells from droplet-based scRNA-seq using whole-genome sequencing variant calls

To match the processing of the whole-genome sequencing datasets, droplet-based sequencing data from decidua and placenta samples were realigned and quantified against the GRCh37 human reference genome using the Cell Ranger Single-Cell Software Suite (v.2.0)^61^. The fetal or maternal origin of each barcoded cell was then determined using the tool demuxlet^67^. Briefly, demuxlet can be used to deconvolve droplet-based scRNA-seq experiments in which cells are pooled from multiple, genetically distinct individuals. Given a set of genotypes corresponding to these individuals, demuxlet infers the most likely genetic identity of each droplet by estimating the likelihood of observing scRNA-seq reads from the droplet overlapping known SNPs. Demuxlet inferred the identities of cells in this study by analyzing each Cell Ranger-aligned BAM file from decidua and placenta in conjunction with a VCF containing the high-quality WGS variant calls from the corresponding mother and fetus (Supplementary Methods). Each droplet was assigned to be maternal, fetal, or unknown in origin (ambiguous or potential doublet), and these identities were then linked with the transcriptome-based cell clustering data to confirm the maternal and fetal identity of each annotated cell type.

### TCR analysis by TraCeR

The TCR sequences for each single T cell were assembled using TraCeR^68^ which allowed the reconstruction of the TCRs from scRNA-seq data and their expression abundance (transcripts per million, TPM), as well as identification of the size, diversity and lineage relation of clonal subpopulations. In total, we obtained the TCR sequences for 1,482 T cells with at least one paired productive αβ or gamma-delta chain. Cells for which more than two recombinants were identified for a particular locus were excluded from further analysis.

### Quantification of KIR gene expression by KIRid

The KIR locus is characterized with high polymorphism both in the number of genes and alleles^9^. Including a single reference sequence for each gene can lead to reference bias for donors that happen to match the reference sequence better. To address these issues, we used a tailored approach in which we mapped each donor’s single cell reads to the corresponding donor-specific reference of KIR alleles, based on existing haplotypes.

### Cell-cell communication analysis

To enable a comprehensive and systematic analysis of the cell-cell communication we have developed CellPhoneDB, a public repository of ligands, receptors and their interactions. Our repository relies on the use of public resources to annotate receptors and ligands (Supplementary methods). We include subunit architecture for both ligands and receptors, in order to accurately represent heteromeric complexes.

Ligand-receptor pairs are defined based on physical protein-protein interaction (PPI) curated by IMEx (http://www.imexconsortium.org/)^69^ and additional extensive manual curation of the literature (Supplementary methods). We provide CellPhoneDB with a user-friendly web interface at www.CellPhoneDB.org/, where the user can search look for ligand/receptor pairs and interrogate their own single-cell transcriptomics data. Membrane protein lists; ligand and receptor lists; and heteromeric complex list are publically available in www.CellPhoneDB.org

To assess the cellular crosstalk between different cell types, we used our repository in a statistical framework for inferring cell-cell communication networks from single cell transcriptome data. We derived potential receptor-ligand interactions based on expression of a receptor by one cell type and a ligand by another cell type, using the high coverage SS2 data. In order to identify the most relevant interactions between cell types, we looked for the cell-type specific interactions between ligands and receptors. Only receptors and ligands expressed in more than 20% of the cells in the specific cluster were considered. First, we randomly permute the cluster labels of each cell 1000 times and determine the mean of the average receptor expression level of a cluster and the average ligand expression level of the interacting cluster. For each receptor-ligand pair this generates a null distribution. By calculating the proportion of the means which are “as or more extreme” than actual mean, we obtain a *p*-value for the likelihood of cell type specificity of a given receptor-ligand pair. We then prioritized interactions that were highly specific based on the number of significant pairs and manually selected biologically relevant ones. For the multi-subunit heteromeric complexes, the member of the complex with the minimum average expression is used for calculating the mean.

### Data and materials availability

The raw sequencing data, expression count data and cell classifications will be deposited at ArrayExpress.

CellPhoneDB and python code can be accessed at cellphonedb.org

## Author contributions

R.V-T. and S.A.T conceived the experiment. Samples were isolated and libraries were prepared by R.V-T with contributions from M.Y.T; J.P; E.S.; S.L. Flow cytometry and FACS experiments were performed by R.V-T; R.A.B; A.F. Immunohistochemistry experiments were performed by M.T; M.E. and R.V-T analysed and interpreted the data with contributions from J.H, A.Z, A.G., M.J.T. Pathological expertise was provided by A.M. R.V-T and S.A.T. wrote the manuscript, with contributions from M.E., K.M, J.H., M.J.T, M.H.. A.M., M.H and S.A.T. co-directed the study. All authors read and accepted the manuscript.

## Acknowledgements

We thank Gerard Graham for the discussions on cytokine interactions; Lira Mamanova and Ayesha Jinat for help on SS2 processing; Ania Hupalowska for help on the illustration; Dorin Popescu and James Fletcher for assistance on tissue processing; Susan Lindsay and Allison Farnworth for ethical help and consenting; Lucy Gardner for help with the immunochemistry experiments; Raghd Rostom and Davis McCarthy for helpful discussions on SNP assignment; Andrew Sharkey for helpful discussions on NK cell biology; Felipe Vieira Braga for helpful discussions on T cell biology; Judith Bulmer for helpful discussions on decidua biology; Tomas Gomes and Ricardo Miragaia for helpful discussions on scRNA-seq analysis; Valentine Svensson and Martin Hemberg for helpful discussions on CellPhoneDB; and all members of the Teichmann and Behjati lab for further discussions on the manuscript. We are indebted to the donors for participating in this research.

Additional funding in support of individual authors was provided as follows: R.V-T is supported by an EMBO Long-Term Fellowship and a Human Frontier Science Program Long-Term Fellowship; J.H. is funded by the Swedish Research Council; S.L. is funded by MRC / Wellcome Trust (#099175/Z/12/Z); M.H. is funded by Wellcome Trust (WT107931/Z/15/Z), The Lister Institute for Preventive Medicine and NIHR and Newcastle-Biomedical Research Centre; S.A.T. is funded by Wellcome Trust, ERC Consolidator and MRG-Grammar.

